# First eight residues of apolipoprotein A-I mediate the C-terminus control of helical bundle unfolding and its lipidation

**DOI:** 10.1101/741546

**Authors:** Gregory Brubaker, Shuhui W. Lorkowski, Kailash Gulshan, Stanley L. Hazen, Valentin Gogonea, Jonathan D. Smith

## Abstract

The crystal structure of a C-terminal deletion of apolipoprotein A-I (apoA1) shows a large helical bundle structure in the amino half of the protein, from residues 8 to 115. Using site directed mutagenesis, guanidine or thermal denaturation, cell free liposome clearance, and cellular ABCA1-mediated cholesterol efflux assays, we demonstrate that apoA1 lipidation can occur when the barrier to this bundle unfolding is lowered. The absence of the C-terminus renders the bundle harder to unfold resulting in loss of apoA1 lipidation that can be reversed by point mutations, such as Trp8Ala, and by truncations as short as 8 residues in the amino terminus, both of which lower the barrier to helical bundle unfolding. Locking the bundle via a disulfide bond leads to loss of apoA1 lipidation. We propose a model in which the C-terminus acts on the N-terminus to destabilize helical bundle. Upon lipid binding to the C-terminus, Trp8 is displaced from its interaction with Phe57, Arg61, Leu64, Val67, Phe71, and Trp72 to destabilize the bundle. However, when the C-terminus is deleted, Trp8 cannot be displaced, the bundle cannot unfold, and apoA1 cannot be lipidated.

## Introduction

Apolipoprotein A-I (apoA1) is the major protein in high density lipoprotein (HDL), and one of the most abundant proteins in human plasma with average levels of ~100 to 150 mg/dl. The assembly of HDL in vivo or by cultured cells is absolutely dependent upon the membrane protein ABCA1, which is defective in Tangier disease (1). However, cell-free reconstituted HDL (rHDL) can be formed from the spontaneous reaction of apoA1 with liposomes made of the short chain phospholipid dimyristoylphosphatidylcholine (DMPC) (2). This reaction has a maximal rate at the DMPC phase transition temperature of ~24°C, where the boundary between the fluid liquid crystalline and gel phases creates lower phospholipid packing density that allows the entry of water and weak detergents (3). The apoA1 protein sequence contains a series of 11 and 22-mer and partial repeats, many of which can form a class A amphipathic alpha helical structure, with a hydrophobic surface bordered by positively charged Lys and Arg residues, and opposed by negatively charged Asp and Glu residues (4). Synthetic class A amphipathic helical peptides such as pA18A, made without sequence similarity to apoA1, can themselves act as weak detergents, and solubilize DMPC liposomes as well as act as ABCA1-dependent acceptors of cellular lipids; however, at high concentrations, some of these synthetic peptides can strip cells of lipids in an ABCA1-independent manner (5). ApoA1, even at high concentrations, does not have the promiscuous ability to accept cell lipids in the absence of ABCA1 (5).

Much about apoA1 function and structure has been learned from studying site-specific substitutions and truncations of apoA1, as well as from various structural studies culminating in the crystal structure of the C-terminal deleted apoA1, solved by Mei and Atkinson in 2011 (6). This crystal structure includes a folded mostly alpha-helical bundle extending from residue 8-112. The apoA1 “consensus” model, built from the crystal structure along with chemical linking, and other biophysical and structural data from the past four decades, has labeled the individual helixes: H1 (residues 8-32), H2 (37-45), H3 (54-64), H4 (68-78), H5 (81-115), and H6 (148-179) (7). It has long been appreciated that the C-terminal truncation of residues 183-243 (called hereafter the ΔC isoform) is dysfunctional in regard to both its DMPC solubilization and ABCA1-dependent lipid acceptor activities (8–11). This was not surprising, as the C-terminus is the most hydrophobic region of apoA1. However, combining the ΔC truncation with deletion of the N-terminal residues 1-43 (called hereafter the ΔN isoform) to create the doubly deleted ΔN/C isoform, completely rescued apoA1’s activity, demonstrating that all that is required for apoA1 function is the central domain from residues 44-182 (10,12). Phillips and colleagues proposed that the C–terminus in full length apoA1 interacts with lipid and transmits a structural change allowing the unfolding of apoA1’s helical bundle revealing its detergent-like activities (13,14). This model and a similar one from Atkinson and colleagues (6,15) can explain why the ΔC is defective in lipid binding, which is recovered by the ΔN/C isoform. In the present study we provide new details on the role of the N-terminal 8 residues and Trp8 in stabilizing the helix bundle and new data on the need for the helical bundle to unfold for apoA1’s lipidation. We support a model that features the central role of Trp8 in maintaining the helical bundle, who’s unfolding is required for apoA1’s lipidation.

## Materials and Methods

### Generation and Purification of recombinant human apoA1 and variants

The bacterial expression vector encoding codon-optimized his-tagged human apoA1 has been previously described (16). All point mutations and deletions were created using the QuickChange II Mutagenesis Kit (Thermo Fisher). All mutations were confirmed by DNA sequencing. Expression plasmids were transformed into *E. coli* BL21 dE3 pLysS and protein expression was induced in shaking cultures by overnight incubation with 0.5 mM Isopropyl β-D-1-thiogalactopyranoside at room temperature. The resulting cellular pellet was resuspended in B-PER lysis solution (Thermo Fisher) containing Lysozyme, DNaseI, and a protein inhibitor cocktail. The cellular debris was removed by centrifugation and the supernatant was diluted into PBS containing 3 M guanidine-HCl. The denatured histidine-tagged apoA1 was purified using Ni Sepharose HP resin (Amersham Biosciences) followed by imidazole elution. Fractions containing recombinant apoA1 were extensively dialyzed against PBS and analyzed for purity by SDS-PAGE and Coomassie Blue staining. Only samples with >95% purity were used.

### Cholesterol Efflux Activity

RAW264.7 cells in 24-well plates were labeled by overnight incubation with DMEM containing 1% fetal bovine serum and 0.5 μCi/mL [^3^H]cholesterol (Perkin Elmer). The next day, the labeling mix was removed and the cells were incubated with DMEM 0.3 mM 8-Br-cAMP for 16 hours to induce endogenous ABCA1 expression. The cells were washed once with DMEM then 5.0 μg/mL apoA1 in 0.5 mL of DMEM + 0.3 mM 8-Br-cAMP was added to each well for incubation at 37°C for 4 hrs. The media was removed and briefly centrifuged and 100 μL was added to a scintillation vial for counting radioactivity to measure of cholesterol released to the media. The cells remaining on the plate were extracted using hexane:isopropanol (3:2, v:v) and the radioactivity was counted as a measure of remaining cellular cholesterol. Percent efflux is calculated as % media counts / (media + cell counts).

### Dimyristoyl phosphatidylcholine (DMPC) liposome clearance assay

DMPC (Avanti Polar Lipids) was dissolved in 2:1 (v:v) chloroform:methanol and dried under nitrogen in glass vials. The lipid was resuspended in PBS by vigorous vortexing and multiple freeze/thaws to prepare multilamellar vesicles (MLVs). The turbid DMPC stock solution was diluted to 0.20 mg/mL in PBS, which yielded an absorbance at 325 nm of < 0.5 AU. ApoA1 samples were tested for their ability to clarify the DMPC vesicles using a Gemini EM microplate reader (Molecular Devices) by adding 250 μL of DMPC to 20 μg of apoAI in PBS (total volume of 300 μL/well). Sample absorbance at 325 nm to assess turbidity was measured over a time course at 24°C.

### 8-Anilino-1-naphthalenesulfonic acid (ANS) Fluorescence Assay

ANS (Sigma Aldrich) at a final concentration of 250 μM was mixed with 30 μg/mL of apoAI samples. Fluorescence spectra were obtained (excitation 395 nm and emission from 425 – 575 nm) using a Gemini EM microplate reader (Molecular Devices). The wavelength of maximal fluorescence (WMF) was determined after data smoothing using GraphPad Prism software.

### Fluorescence spectroscopy and guanidine or thermal denaturation

For guanidine denaturation, apoA1 samples were prepared at 10 μg/mL in increasing amounts of guanidine hydrochloride (0, 0.25, 0.5, 0.75, 1.0, 1.25, 1.5, 1.75, 2.0, 2.25, 2.5 M) at room temperature. For thermal denaturation 10 μg/mL apoA1 in PBS was equilibrated at the indicated temperature. For both assays, fluorescence spectra were measured using an excitation of 295 nm and emission from 325 – 375 nm using a Gemini EM microplate reader (Molecular Devices) at the temperature of equilibration. The WMF was determined after data smoothing using GraphPad Prism software. A red shift is observed as the environment of the four apoA1 tryptophans goes from a hydrophobic to an aqueous environment during the unfolding process.

### ApoA1 structural modeling

The ΔC apoA1 crystal structure (Protein Data Bank accession number 3r2p) (6) and the full length apoA1 consensus model (downloaded from htpp://homepages.uc.edu/~davidswm/structures.html) (7) were visualized and customized using PyMOL 2.1.1 (Incentive Product).

### Statistical analysis

All data show mean + SD. Cell well and sample replicates were used and assumed to be parametrically distributed, and as our coefficients of variation were small and no in vivo measures were ascertained. Multiple columns of data were compared by one-way ANOVA with Dunnett’s posttest against one control column as indicated in the figure legends. Statistics were performed using GraphPad Prism software.

## Results

### Central Domain of apoA1 is sufficient for apoA1’s cell-free and cellular lipidation

N-terminal his-tagged recombinant human apoA1 (rh-apoA1) was purified for the full length wild type (WT) isoform, along with the N-terminal residues 1-43 deleted (ΔN), the C-terminal residues 185-243 deleted (ΔC), and the N- and C-terminal double deleted (ΔN/C) isoforms, the latter of which contains only the central domain of apoA1. We confirmed prior studies that the WT, ΔN, and ΔN/C isoforms were competent to clear DMPC MLVs, while the ΔC isoform was incapable of clearing these liposomes (Figure 1A) (10,12). This same pattern of activity was also observed for ABCA1-mediated cholesterol efflux from RAW264.7 murine macrophages (Figure 1B).

**Fig 1.**
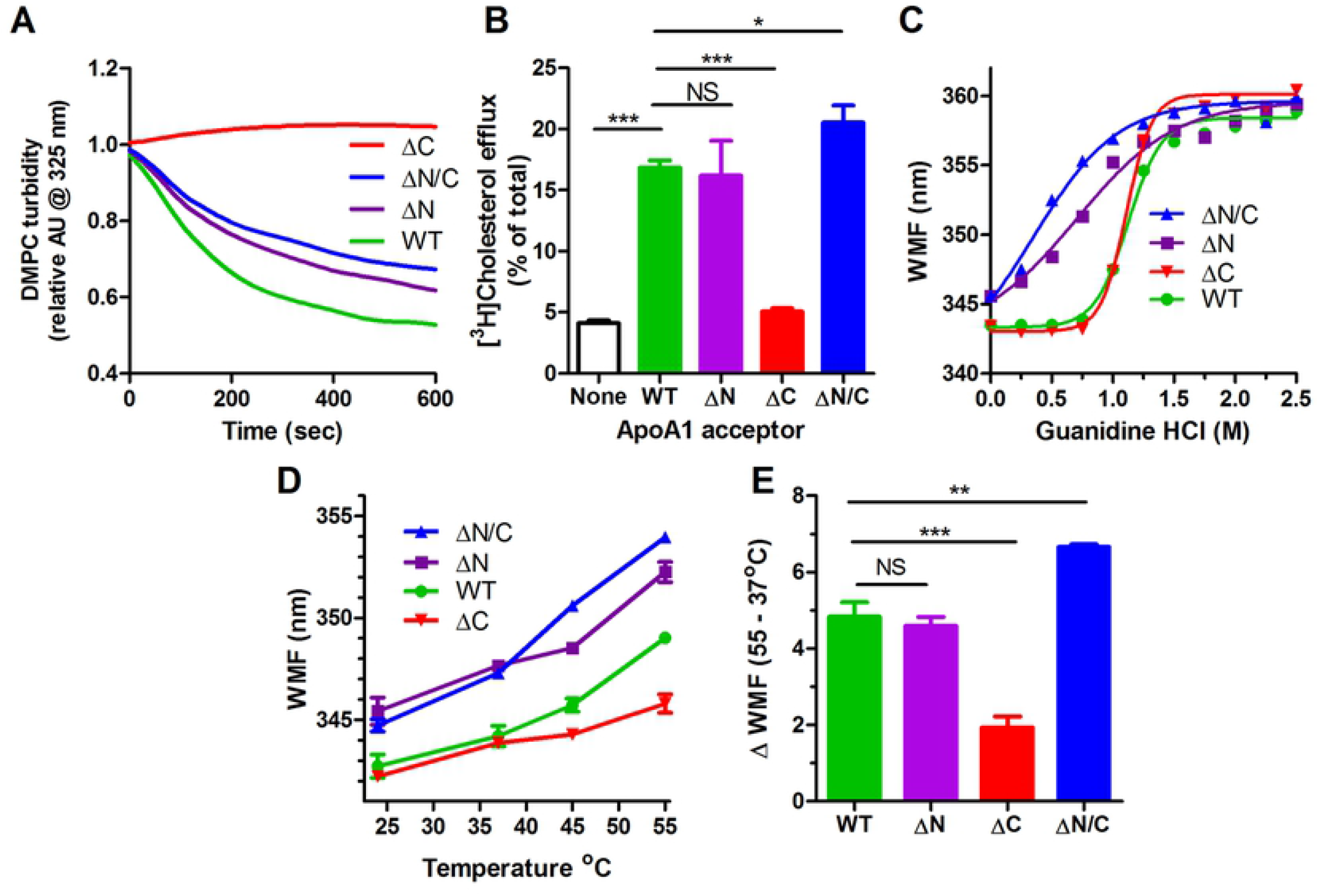
ApoA1 ΔN/C double deletion restores activity to ΔC deletion. **A.** DMPC MLV clearance by apoA1. Each line is the average of triplicate wells. Different apoA1 isoforms: green, wildtype (WT); purple, ΔN; red, ΔC; blue, ΔN/C. **B.** Cholesterol efflux from ABCA1-induced RAW264.7 cells to different apoA1 isoforms. **C.** Guanidine denaturation of apoA1 isoforms assayed by wavelength of maximal fluorescence (WMF) of endogenous Trp residues. **D.** Thermal denaturation of apoA1 isoforms assayed by Trp WMF in triplicate wells. **E.** Thermal denaturation of apoA1 isoforms as the delta WMF between 55 and 37°C. The bar graphs are mean + SD, n=3 (***, p<0.001; **, p<0.01; *, p<0.05; NS, not significant; by one way ANOVA with Dunnett’s posttest compared to WT apoA1).

### ApoA1 helical bundle unfolding of different isoforms

The crystal structure of the lipid-free ΔC apoA1 isoform was determined by Mei and Atkinson (6), revealing a large alpha-helical bundle composed of three segments: segment 1, residues 8-40; segment 2, residues 41-67 that are antiparallel to segment 1; and segment 3, residues 68-115 that are parallel to segment 1). We hypothesized that the apoA1 C-terminal domain can assist in destabilizing the structure of this bundle, and that the unfolding of the bundle is required for apoA1’s lipidation. To examine the ability of the various apoA1 isoforms to unfold, we performed guanidine denaturation dose response studies, where unfolding was monitored by the WMF red shift in endogenous Trp fluorescence as it moves from a hydrophobic to hydrophilic environment (9). There are four Trp residues in apoA1 at positions 8, 50, 72, and 108, all of which are found in the helical bundle, although, Trp 8 is just at the boundary. The WMF assay is not dependent upon the concentration of apoA1, and thus is insensitive to small changes in concentration among the four apoA1 isoforms. The EC50 for unfolding of the WT and ΔC isoforms was similar at 1.12 and 1.11 M guanidine, respectively (Figure 1C). The ΔN isoform, missing the anchor residues for the segment 1 and the beginning of segment 2 of the helical bundle, unfolded much more readily with an EC50 of 0.70 M guanidine, while the ΔN/C isoform unfolded even more easily with an EC50 of 0.35 M guanidine (Figure 1C). The two N-terminal deletion isoforms (ΔN and ΔN/C) had much lower Hill slopes, indicating less cooperativity in unfolding, vs. the other two isoforms. In addition, in the absence of guanidine, both N-terminal deletion isoforms have higher basal WMFs than the WT and ΔC isoforms. We performed thermal denaturation in the absence of guanidine and again found that the two N-terminal deletion isoforms (ΔN and ΔN/C) unfolded better (larger WMF shift) than the WT and ΔC isoforms (Figure 1D), which also confirmed the baseline red shift of the two N-terminal deletion isoforms at 24°C. Thermal denaturation showed that the WT isoform was slightly easier to unfold than the ΔC isoform, thus showing that the C-terminus stabilized the folded state (Figure 1E).

### Role of Trp8 and the N-terminal residues in helical bundle unfolding and apoA1 lipidation

The baseline red shift of the N-terminal deletion isoforms could represent either a more unfolded state of the helical bundle or merely the loss of the fluorescence signal from Trp8, which is missing in these isoforms. To evaluate this, we mutated Trp8 to Phe (W8F), Leu (W8L), or Ala (W8A) and we found that the W8F and W8L, two bulky hydrophobic substitutions, retained the baseline WMF of the WT isoform, while smaller W8A substitution increased the baseline WMF (Figure 2A). Subjecting these isoforms to guanidine denaturation showed a stepwise pattern going from least to most sensitive to guanidine unfolding for the aromatic to larger hydrophobic to smaller hydrophobic residues at position 8 (Figure 2A, guanidine EC50: WT, 1.11M; W8F, 0.83M; W8L, 0.77M; W8A, 0.70M).

**Fig 2.**
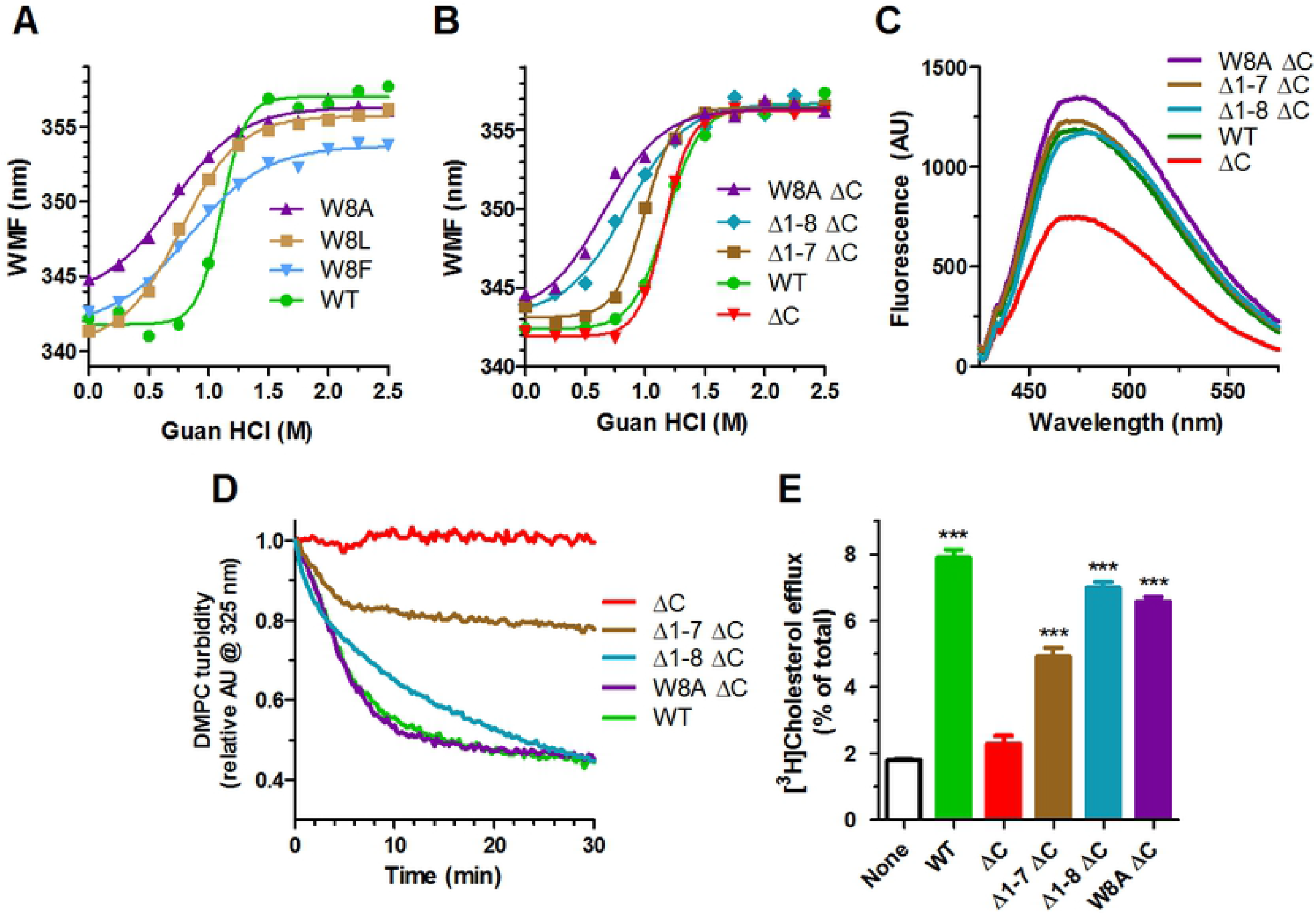
Role of Trp8 apoA1 helix bundle unfolding and rescue of ΔC isoform activity. **A.** Guanidine denaturation of apoA1 isoforms assayed by Trp WMF. Different apoA1 isoforms: green circles, wildtype (WT); blue inverted triangles, W8F; brown squares, W8L; purple triangles, W8A. **B.** Guanidine denaturation of apoA1 isoforms assayed by Trp WMF. Different apoA1 isoforms: green circles, wildtype (WT); red inverted triangles, ΔC; brown squares, Δ1-7 ΔC; blue diamonds, Δ1-8 ΔC; purple triangles, W8A ΔC. **C.** Hydrophobicity of apoA1 isoforms assayed by ANS fluorescence, same colors as in panel B. **D.** DMPC MLV clearance by apoA1, same colors as in panel B **E.** Cholesterol efflux from ABCA1-induced RAW264.7 cells to different apoA1 isoforms (average + SD; ***, p<0.001 by one way ANOVA with Dunnett’s posttest compared to no apoA1).

To further examine the role of Trp8, we made the least stable point mutation W8A, and two new N-terminal deletions 1-7 (Δ1-7) and 1-8 (Δ1-8), all on the ΔC background. The first 7 residues contain 3 prolines in positions 3, 4, and 7 Guanidine denaturation demonstrated that the Δ1-8 and W8A were most sensitive to unfolding, while the Δ1-7 had intermediate sensitivity (Figure 2B). The unfolding sensitivity completely aligned with the DMPC clearance and ABCA1-mediated acceptor activities of these ΔC isoforms, such that the easiest to unfold isoforms (W8A and Δ1-8) completed rescued the ΔC loss of function, while the Δ1-7 only partially rescued these activities (Figure 2C, D, E). Thus, Trp8 has an essential role in stabilizing the helical bundle, as the least conservative substitution, W8A, or deletion of the first 8 residues, destabilize the bundle and allow for functional recovery of the ΔC isoform.

### Locking apoA1’s helical bundle diminished its cell-free and cellular lipidation

To test the role of unfolding of the helical bundle on apoA1’s lipidation, we used site directed mutagenesis to replace residues Leu38 and Met112, which in the crystal structure (6) are predicted to be only 3.4 angstroms apart, with Cys residues to create the 38C/112C helix bundle locked apoA1 isoform (Figure 3A). After purification, non-reducing SDS PAGE revealed that all of the disulfide bonds are intra-molecular, as we did not observe any dimer sized bands (Figure 3B). The 38C/112C locked apoA1 isoform was harder to unfold in guanidine (EC50 = 1.56 M guanidine) vs. the WT isoform; but, it regained sensitivity to guanidine unfolding upon unlocking the N-hairpin by disulfide reduction with DTT (Figure 3C). The 38C/112C locked apoA1 isoform had very little DMPC MLV clearance activity compared to WT apoA1; however, upon unlocking the helix bundle with DTT, this activity was largely restored, indicating that loss of this activity was due to the locked helix bundle rather than to the double Cys substitution (Figure 3D). The cellular ABCA1-mediated efflux capacity of the 38C/112C locked apoA1 isoform was reduced by 75% vs. the WT isoform; however, this activity was completely restored by reductive methylation of the disulfide bond using N-ethylmaleimide (NEM) to unlock the helix bundle (Figure 3E). These studies demonstrate that unfolding the helical bundle is necessary in order for apoA1 lipidation via solubilization of DMPC vesicles or via cellular ABCA1 activity.

**Fig 3.**
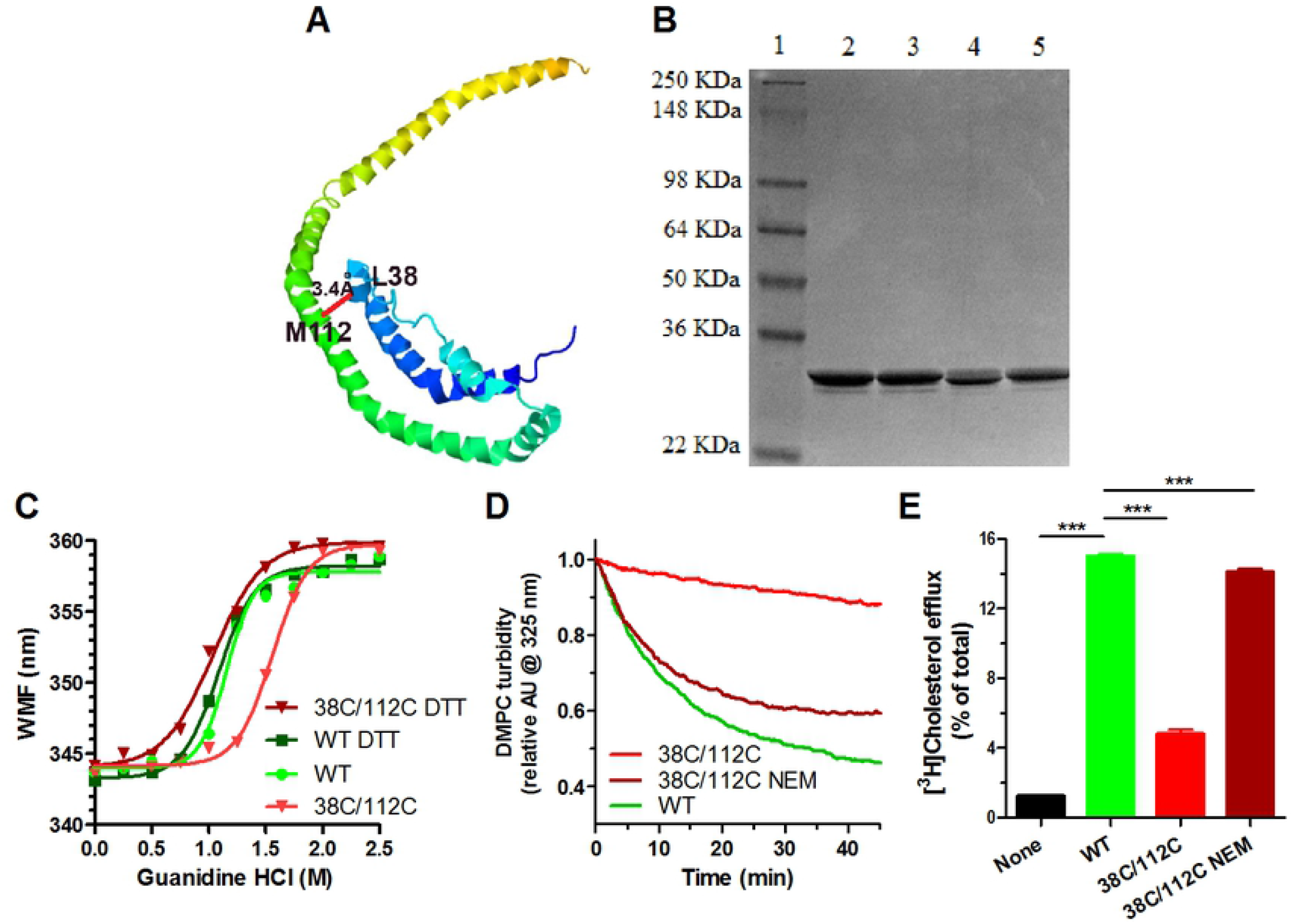
Disulfide lock of the helical bundle impedes apoA1 activity. **A.** Portion of the crystal structure of ΔC apoA1 from (6), showing proximity of L38 and M112 at the “top” of the helical bundle. Color scheme shows N-terminus with dark blue towards the C-terminus in orange. **B.** Coomassie blue stained SDS PAGE of purified recombinant apoA1 isoforms, showing only intramolecular disulfide bonds in the 38C/112C isoform. Lane 1, MW marker; lane 2, WT w/o DTT; lane 3 WT + DTT; lane 4, 38C/112C w/o DTT; lane 5, 38C/112C + DTT. **C.** Guanidine denaturation of apoA1 isoforms assayed by Trp WMF. Different apoA1 isoforms: light green circles, non-reduced WT; dark green squares WT + DTT; inverted bright red triangles, nonreduced 38C/112C; inverted dark red triangles, 38C/112C + DTT. **D.** DMPC MLV clearance by apoA1; green, WT; bright red, non-reduced 38C/112C; dark red, 38C/112C reductively methylated with NEM. **E.** Cholesterol efflux from ABCA1-induced RAW264.7 cells to different apoA1 isoforms (average + SD; ***, p<0.001 by one way ANOVA with Dunnett’s posttest compared to WT apoA1).

### Proline substitutions in the helix bundle rescue the activity of Δ185-243 apoA1

To further prove that it is the ability of the helical bundle to unfold that can rescue the activities of the ΔC isoform, we replicated the study of Tanaka et al. (17) and substituted Tyr18 and Ser55, within the N-terminal helix hairpin bundle, with proline residues on the full length and ΔC background isoforms, creating the 2P isoforms. The 2P-WT and 2P-ΔC isoforms are much less folded as they have a large red-shifted baseline WMF, and they are much more sensitive to guanidine unfolding compared to the WT and ΔC isoforms (Figure 4A). As previously shown (17), the 2P-ΔC isoform completely restored to WT levels the impaired activities of the ΔC isoform in regard to both DMPC clearance and ABCA1-mediated cholesterol efflux (Figure 4B, C). Thus, like the Trp8, D18, and ΔN/C isoforms, these specific proline substitutions promote the helical bundle unfolding and rescue the activity deficits of the ΔC isoform, indicating that the C-terminus is dispensable for apoA1 lipidation when the helical bundle is destabilized.

**Fig 4.**
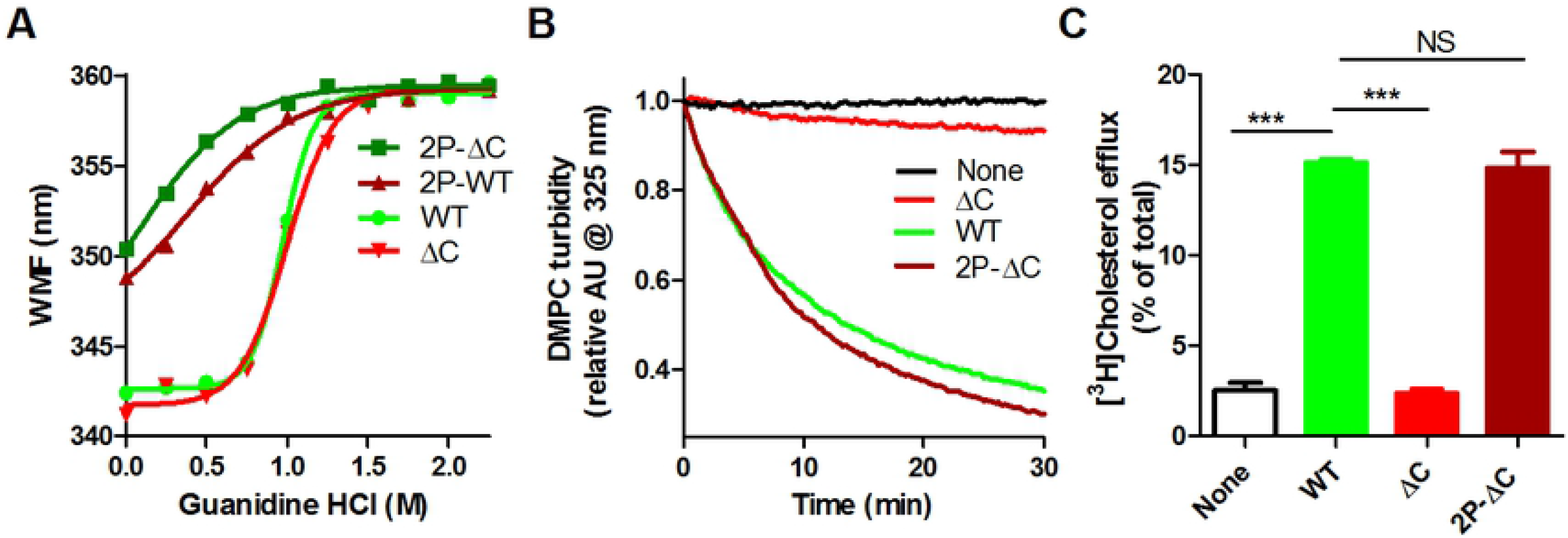
ApoA1 with two proline substitution in helical bundle restores activity of the ΔC isoform. **A.** Guanidine denaturation of apoA1 isoforms assayed by Trp WMF. Different apoA1 isoforms: light green circles, non-reduced WT; dark green squares 18P/55P (2P-WT); inverted bright red triangles, ΔC; dark red triangles, 2P-ΔC. **B.** DMPC MLV clearance by apoA1; same colors as in panel B. **C.** Cholesterol efflux from ABCA1-induced RAW264.7 cells to different apoA1 isoforms, same colors as in panel A (average + SD; ***, p<0.001 by one way ANOVA with Dunnett’s posttest compared to WT apoA1).

## Discussion

We propose a model for regulation of apoA1’s helix bundle unfolding based on the ΔC isoform crystal structure (6), the “consensus” model of apoA1 (7), and our current findings. The crystal structure places Trp8 (W8) at the start of H1 (6), which we propose to be a central player that acts as a “node” to stabilize the helix bundle. W8 sits in a pocket flanked by multiple residues within H3, H4, and the linker region between H3 and H4 (Figure 5). W8 may have hydrophobic interactions with residues F57 (4.1Å distance from W8 in H3), L64 (4.0Å in H3), V67 (4.0 Å in the linker), F71 (5.3 Å in H4) and W72 (3.7 Å in H4). In addition, W8 is reported to have a pi-cation interaction with R61 (3.6 Å in H3) (6), which combined with the above hydrophobic interactions are reported to be the “major forces that hold the helical bundle together (6). Thus, the Trp8 substitution W8A or deletion of residues 1-8 removes the “node” and allows the helical bundle to unfold easier.

**Fig 5.**
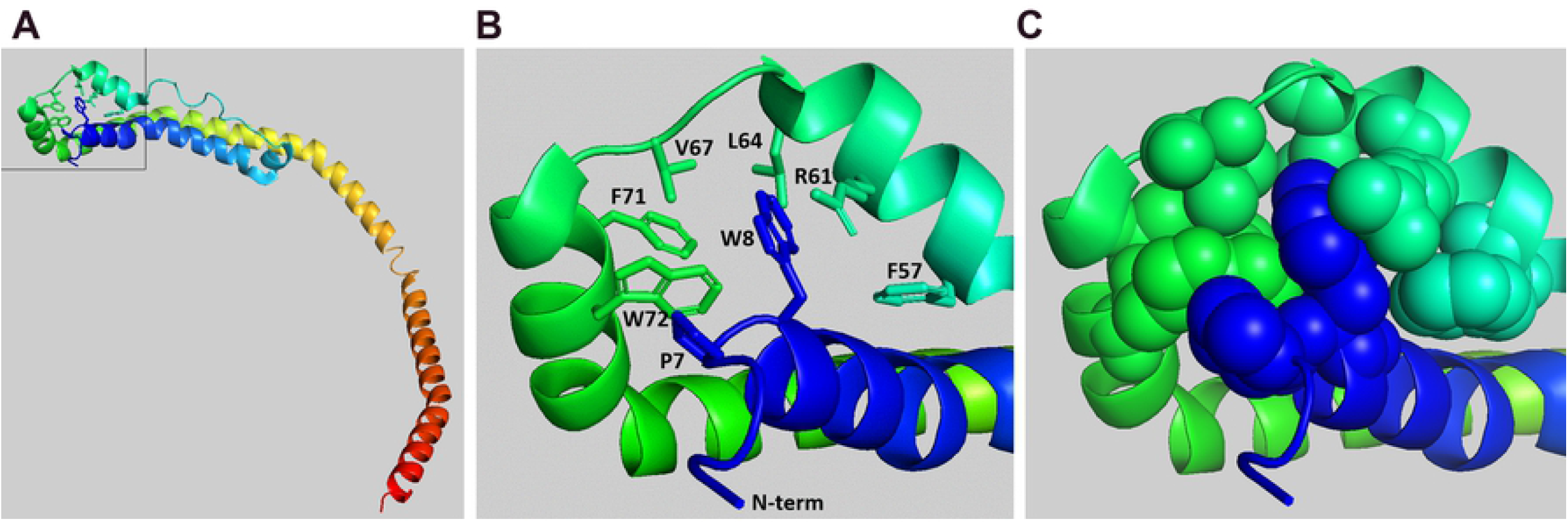
Trp8 in helix 1 coordinates interaction with residues in helices 2 and 3. **A.** Crystal structure of the ΔC isoform from (6) with N-terminus in blue and C-terminus in red. Boxed area is shown in subsequent panels. **B.** Ribbon and stick diagram showing Trp8 and the residues in close proximity in helices 2 and 3 as indicated. **C.** Ribbon and sphere diagram showing Trp8 and the residues indicated in panel B.

In the apoA1 structure consensus model, the C-terminus is adjacent to the N-terminus (Figure 6A, B) (7). This interaction is supported by numerous lysine/amine crosslinking studies using full length, monomeric, lipid-free apoA1, including cross links between the following lysine pairs (first lysine/amine in the N-terminus/hairpin bundle between residues 1 and 115 x second lysine in the C-terminus between residues 185 and 243): 1 x 195; 1 x 208; 1 x 239; 12 x 195; 12 x 226; 12 x 239; 23 x 239; 40 x 239; 58 x 239; 59 x 195; 59 x 239; 77 x 195; 77 x 208; 77 x 239; 87 x 239; 88 x 195; 94 x 226;94 x 239;96 x 195, 96 x 208; 96 x 226; 96 x 239; 107 x 208, and 107 x 239 (7,18–20). Biophysical studies have previously shown that the hydrophobic C-terminus can undergo a transition to acquire more alpha-helical structure upon lipid binding (21). We propose that the newly formed amphipathic alpha helical C-terminus tugs on the N-terminal 7-residue appendage to pull W8 out of its pocket (Figure 6C, D). This pulling would disrupt W8’s interactions with F57, R61, L64, V67, V69, F71, and F72, allowing the helix bundle to unfold so that the newly exposed amphipathic helical hydrophobic surface may bind lipid for HDL assembly. When the C-terminus is deleted W8 cannot be pulled out from its helix bundle stabilizing position, preventing bundle unfolding and apoA1 lipidation. The bundle destabilizing variants that remove the W8 node (W8A, Δ1-8) at the base of the helix bundle or otherwise destabilize the helical bundle (W18P, S55P in the middle of the helix bundle) can completely rescue the activity of the ΔC isoform. Similarly, the N-terminal 7 residues may also help to stabilize the position of W8, with P7 4.2 Å away from W72, such that the Δ1-7 isoform partially rescues the activity of the ΔC isoform.

**Fig 6.**
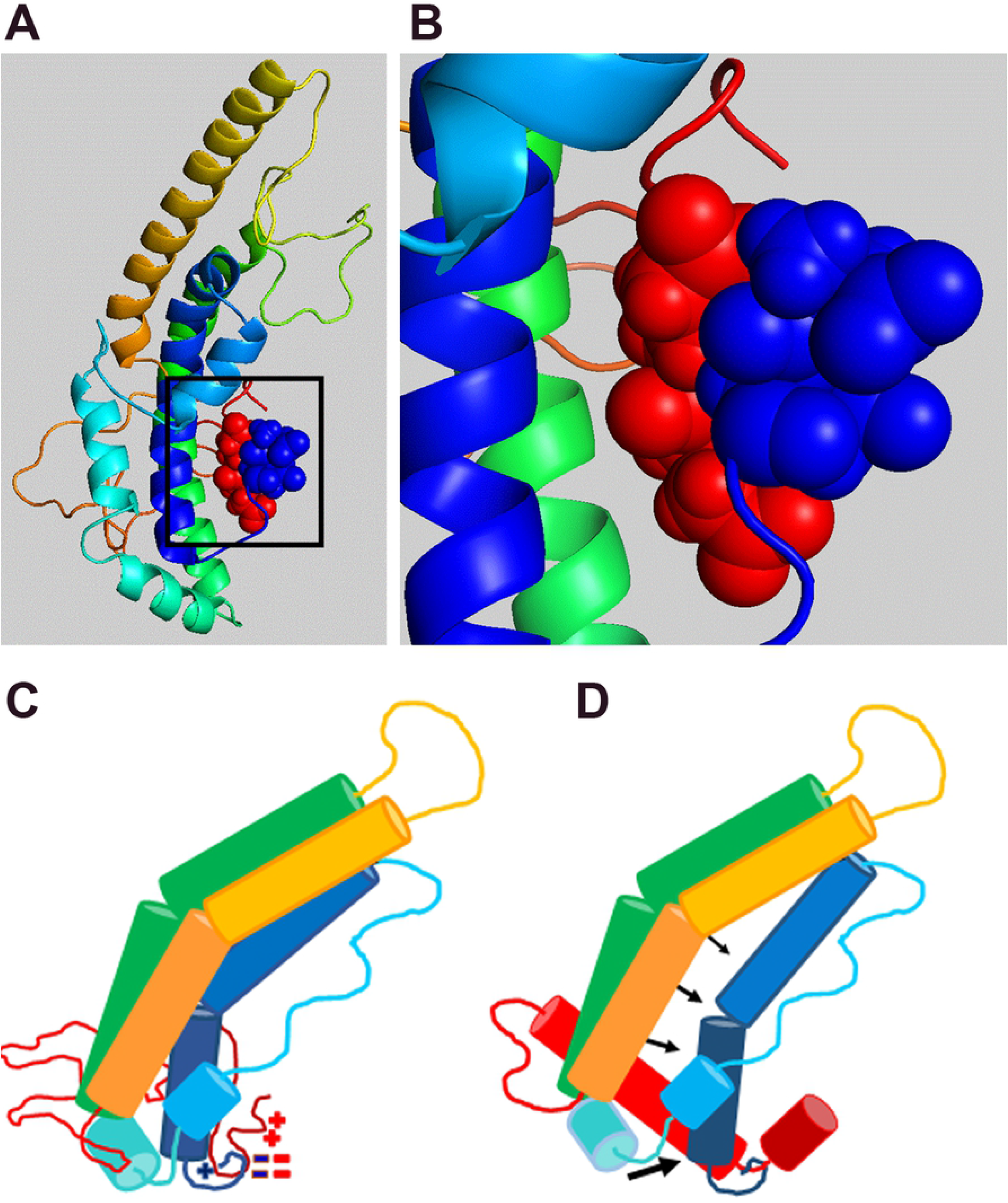
Model for N- and C-terminal interaction to open helical bundle upon C-terminus lipid sensing. **A.** Consensus model of monomeric lipid-free apoA1 from (7) with N-terminus in blue and C-terminus in red, showing close proximity of N-and C-termini. **B.** Blowup of boxed region panel A showing proximity of surfaces for residues D1, E2, E234, and E235. **C.** Cartoon version of ΔC crystal structure from (6) fused to the unstructured C-term region from (7) showing helical segments and proximity of charged residues at N- and C-termini. **D.** Cartoon showing C-terminal helical transformation and extension upon lipid sensing, leading to displacement of the N-terminus. This results in pulling Trp8 out of its position coordinating helices 1 through 3, allowing the unfolding of the helical bundle to expose apoA1’s detergent-like amphipathic helices that is required for its lipidation.

Another helical bundle destabilizing isoform (L38G, K40G), in the context of full length apoA1, was also shown to unzip the helical bundle from the opposite end of W8, exposing more hydrophobic surface and leading to faster cellular ABCA1 mediated HDL formation (22). However, whether this bundle destabilizing isoform could rescue the activity of C-terminal deletion was not determined (22).

Philips and colleagues first proposed that the C-terminal domain senses lipids leading to the opening of apoA1’s N-term helix bundle allowing the exposure of its hydrophobic surfaces and its subsequent lipidation (13,14). This model has been supported by electron paramagnetic resonance evidence showing a structural change in the C-terminal domain upon lipid binding (21), which may drive subsequent unfolding of the helical bundle. Atkinson and colleagues x-ray crystal and apoA1 site directed mutagenesis data support the model that N-terminal helical bundle unfolding is required for lipidation (6,22). Our current findings extend the evidence for this model by showing that locking the helical bundle with a disulfide bond inhibits apoA1 lipidation, and that W8 plays a central role in coordinating one end of the helical bundle.

All four Trp residues in apoA1 occur in the helical bundle (residues 8, 50, 72, and 108). Our prior in vitro mutagenesis work showed that Trp oxidation is responsible for the myeloperoxidase induced loss of apoA1 DMPC solubilization and ABCA1-mediated cholesterol acceptor activities, as the 4WF isoform, with all 4 Trp residues replaced by non-oxidizable Phe residues, is resistant to loss of these activities (23). Substitution of just Trp72 with Phe (W72F) protects apoA1 from MPO induced loss of activity by ~50%, with the other three Trp to Phe substitutions residues responsible for the other 50% of protection (24). Thus, the oxidation of the helical bundle Trp residues may either prevent the unfolding of the helical bundle, or alternatively disrupt the amphipathic alpha helical surface required for apoA1 lipidation.

ApoA1 is one of the most abundant plasma proteins with normal levels of ~ 1.5 mg/ml. We hypothesize that apoA1 evolved with its helical bundle structure in order to protect cells from a promiscuous detergent like activity of an extended amphipathic helix. Remaley has previously shown that high concentrations of amphipathic helical peptides can induce lipid efflux from cells even in the absence of ABCA1 expression; while apoA1 induced lipid efflux is dependent upon ABCA1 expression (5). We demonstrated that ABCA1 mediates the flop of both phosphatidylinositol 4,5 bisphosphate and phosphatidylserine, the former of which is required for apoA1 binding to ABCA1 expressing cells, and the latter of which increases cholesterol extractability to apoA1 or other weak detergents (25). We further showed that apoA1 binding to ABCA1 expressing cells leads to unfolding of the helical bundle by the use of self-quenching fluorescent probes at positions 38 and 112 (26). Thus, it appears that apoA1 and ABCA1 co-evolved to regulate apoA1 helical bundle unfolding in order to tightly regulate the detergent-like activity of the abundant plasma protein apoA1.

